# The Epstein-Barr Virus miR-BHRF1 microRNAs Regulate Viral Gene Expression in *cis*

**DOI:** 10.1101/170746

**Authors:** Brigid Chiyoko Poling, Alexander M. Price, Micah A. Luftig, Bryan R. Cullen

## Abstract

The Epstein-Barr virus (EBV) miR-BHRF1 microRNA (miRNA) cluster has been shown to facilitate B-cell transformation and promote the rapid growth of the resultant lymphoblastoid cell lines (LCLs). However, we find that expression of physiological levels of the miR-BHRF1 miRNAs in LCLs transformed with a miR-BHRF1 null mutant (∆123) fails to increase their growth rate. We demonstrate that the pri-miR-BHRF1-2 and 1–3 stem-loops are present in the 3’UTR of transcripts encoding EBNA-LP and that excision of pre-miR-BHRF1-2 and 1–3 by Drosha destabilizes these mRNAs and reduces expression of the encoded protein. Therefore, mutational inactivation of primiR-BHRF1-2 and 1–3 in the ∆123 mutant upregulates the expression of not only EBNA-LP but also EBNA-LP-regulated mRNAs and proteins, including LMP1. We hypothesize that this overexpression causes the reduced transformation capacity of the ∆123 EBV mutant. Thus, in addition to regulating cellular mRNAs in *trans*, miR-BHRF1-2 and 1–3 also regulate EBNA-LP mRNA expression in *cis*.

**Highlights:**

- The EBV miR-BHRF1 microRNAs do not up upregulate B cell growth in *trans*.
- EBNA-LP expression is downregulated by pri-miR-BHRF1-2 and 1–3 acting in *cis*.
- Loss of miR-BHRF1-2 and 1–3 causes EBNA-LP overexpression and inhibits B cell growth.
- Novel alternative splicing of EBV Cp/Wp transcripts was identified.

## Introduction

Epstein Barr Virus (EBV) is a double-stranded DNA virus of the γ-herpes subfamily that infects >90% of the global population. EBV is the causative agent of various malignancies including diffuse large B cell lymphoma (DLBCL), Burkitt lymphoma, post-transplant lymphoproliferative disorder (PTLD) and nasopharyngeal carcinoma. Additionally, EBV infection of naïve young adults results in infectious mononucleosis. During primary infection, EBV infects resting B cells and establishes a latency stage III infection, which is associated with DLBCL and PTLD. During latency III, EBV-infected B cells express the full repertoire of latency proteins (EBNA-LP, EBNA1, 2, 3A, 3B, 3C, LMP1 and 2A/B) as well as non-coding RNAs, including the EBV-encoded small RNAs (EBERs) and several microRNAs (miRNAs). In the laboratory, EBV latency III can be modeled by infection of primary human B cells to generate lymphoblastoid cell lines (LCLs) (Rickinson and Kieff, 2007).

EBNA-LP and EBNA2 are the first viral proteins expressed after B cell infection, through activation of the W promoter (Wp) by B cell specific transcription factors (Bell et al., 1998; Kirby et al., 2000; Tierney et al., 2000). EBNA2 and EBNA-LP then activate the Cp promoter, to drive expression of EBNA-LP, EBNA-1, 2, 3A, 3B and 3C, as well as the LMP1 promoter (LMP1p) (Harada and Kieff, 1997; Nitsche et al., 1997; Peng et al., 2005; Sung et al., 1991; Woisetschlaeger et al., 1990). Because the viral mRNAs encoding the EBNA proteins are all transcribed from the same promoters, alternative splicing is critical for their appropriate expression. EBNA-LP is the first open reading frame (ORF) after both the Cp and Wp promoters and EBNA-LP expression is dependent on a complex pattern of alternative splicing. First, EBNALP mRNAs require splicing from the first exon of the Wp transcripts (W0) early after infection, or the second exon of the Cp transcripts (C2) late in infection, to provide the initial AT dinucleotide in the ATG codon that initiates EBNA-LP translation. Second, EBNA-LP mRNAs must splice internally into the W1 exon, at the W1’ 3’ splice site, to incorporate the G in the ATG. EBNA-LP mRNAs then continue splicing to integrate multiple copies of the W1/W2 exon repeats in frame, which results in EBNA-LP proteins with a range of different molecular weights. The B95-8 strain of EBV encodes 11 W1/W2 repeats and it was previously shown that incorporation of a minimum of five repeats is necessary for full EBNA-LP function (Tierney et al., 2011). EBNA-LP mRNAs then splice from the terminal W2 exon to the Y1 and Y2 exons, where the translational stop codon for EBNA-LP is located in the Y2 exon. Y2 can then be spliced to several different 3’ exons, including those that encode EBNA1, 2, 3A, 3B, or 3C. If the ATG initiation codon for EBNA-LP is not generated by appropriate splicing at the 5’ end of these viral transcripts, then a long 5’UTR for the mRNAs encoding the 3’ EBNA1, 2, 3A, 3B, or 3C open reading frames (ORF) is generated instead (Rogers et al., 1990).

In addition to proteins, EBV encodes two miRNA clusters named for their location in the EBV genome. The BamH1 fragment H rightward facing 1 (BHRF1) miRNA cluster consists of three pri-miRNA stem-loops, generating the four mature miR-BHRF1-1, 1–2, 1–2*, and 1–3 miRNAs (miR-BHRF1-2 gives rise to almost equal levels of two miRNAs derived from each arm of the pre-miRNA intermediate). The miR-BHRF1-1 miRNA is located in the promoter for the late BHRF1 mRNA while miR-BHRF1-2 and 1–3 are located in the BHRF1 mRNA 3’ UTR. Additionally, the BamH1 fragment A rightward transcript (miR-BART) miRNA cluster is composed of 22 primiRNAs expressed from the various BART transcripts (Cai et al., 2006; Grundhoff et al., 2006; Pfeffer, 2004; Zhu et al., 2009). The miR-BHRF1 miRNAs are only expressed during latency III while the miR-BART miRNAs are expressed during all stages of latency, though at reduced levels in latency III (Cai et al., 2006; Zhu et al., 2009).

While the EBV-encoded miRNAs were discovered over a decade ago, and photoactivatable ribonucleoside-enhanced crosslinking and immunoprecipitation (PAR-CLIP) has allowed for the identification of many of their mRNA target sites (Riley et al., 2012; Skalsky et al., 2012), elucidating their role during EBV-infection has remained challenging. So far, EBV-encoded miRNAs have been shown to play a role in immune evasion (Albanese et al., 2016; Tagawa et al., 2016; Xia et al., 2008), apoptosis (Kang et al., 2015; Lei et al., 2013), and inhibition of tumor suppressors (Bernhardt et al., 2016). Additionally, inactivation of the miR-BHRF1 miRNA cluster results in a reproducible and robust reduction in B cell transformation and LCL growth (Feederle et al., 2011a, 2011b; Haar et al., 2015; Seto et al., 2010), which suggested that the miR-BHRF1 miRNA cluster was promoting EBV transformation by downregulating the expression in *trans* of cellular mRNAs that inhibit this process. We tested this hypothesis by ectopically expressing the miR-BHRF1 miRNAs in LCLs generated using a previously described EBV mutant that lacks all these viral miRNAs (∆123 LCLs). However, we were unable to rescue the slow growth rate of ∆123 LCLs, when compared to WT LCLs, even when all the miR-BHRF1 miRNAs were simultaneously expressed at or above physiological levels. Furthermore, we observed that the ∆123 LCLs expressed a greatly increased level of the EBV transcription factor EBNA-LP when compared to WT LCLs derived from the same donor. Here, we present evidence in favor of the alternative hypothesis that the miR-BHRF1 miRNAs are in fact acting in *cis* to downregulate the expression of EBNA-LP to a level optimum for B cell transformation by inducing the cleavage of EBNA-LP mRNAs during miRNA excision.

## Results

### Trans-complementation does not rescue the growth of LCLs lacking the miR-BHRF1 microRNAs

Previous studies implicated the miR-BHRF1 miRNAs in enhancing the transformation of primary B cells into LCLs, as well as increasing their cell cycle progression, and this enhanced growth rate was maintained in established LCLs (Feederle et al., 2011a, 2011b; Seto et al., 2010). Our original goal was to identify the individual miR-BHRF1 miRNAs responsible for efficient growth of wildtype (WT) LCLs and we therefore first confirmed the slow growth phenotype of ∆123 LCLs. As shown in Fig. 1A, we indeed observed that WT LCLs grow approximately twice as fast as ∆123 LCLs derived from the same blood donor.

**Figure 1.**
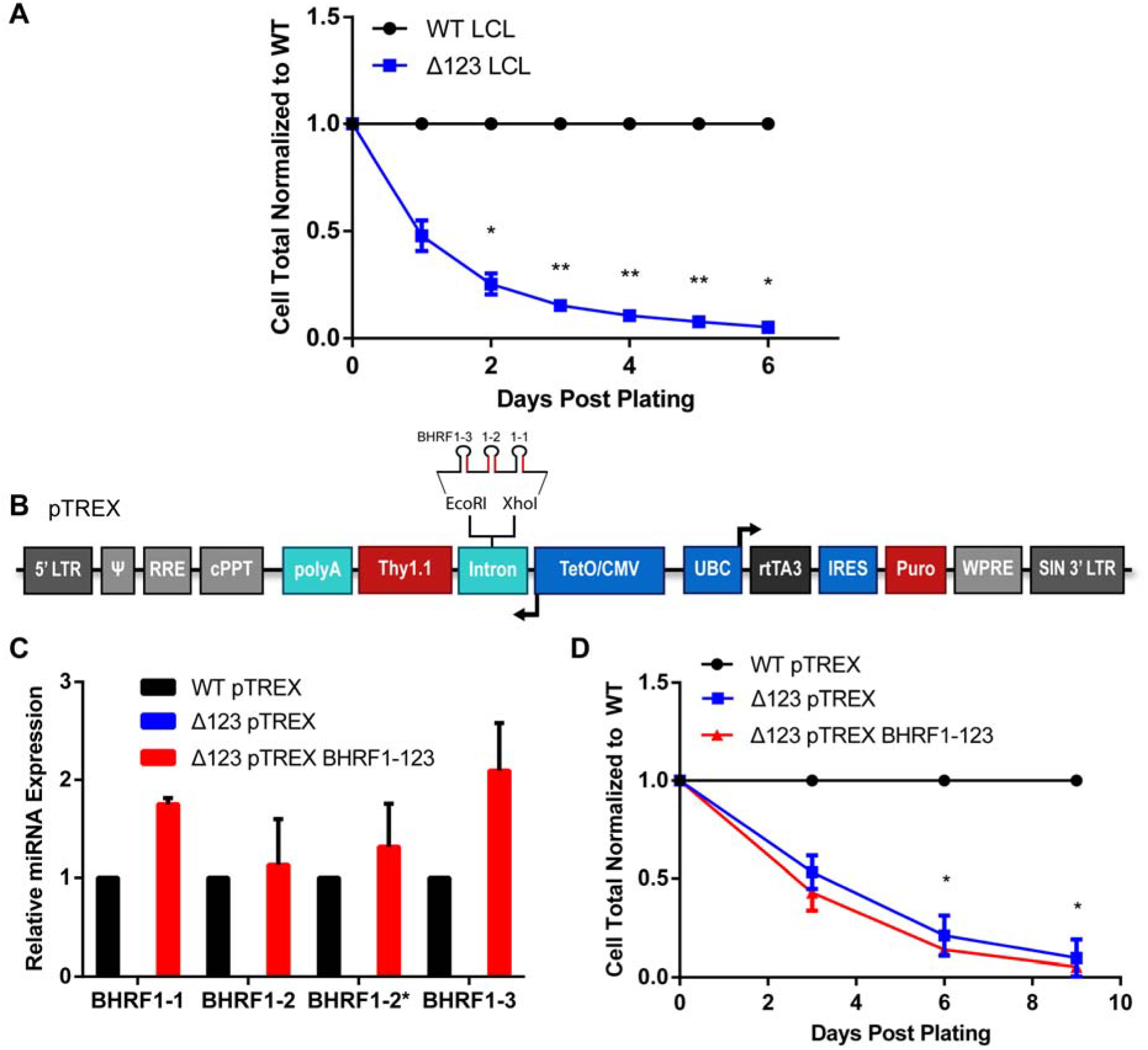
Ectopic expression of the BHRF1 miRNAs does not rescue rapid ∆123 LCL growth. (A) Total intact cells were counted by flow cytometry for WT and ∆123 LCLs every day for six days. ∆123 LCL total cell count was normalized to WT cell total, which was set at 1.0. Average of three donors. *p<0.05, **p<0.01; one-sample t-test. (B) Schematic of the pTREX vector used to express the BHRF1 miRNAs (Poling et al., 2017). Pri-miRNAs, shown in red, were cloned between the XhoI and EcoRI sites present inside the intron 5’ of Thy1.1. miRNA expression was driven from an antisense tet-inducible minimal CMV promoter (TetO/CMV). pTREX also contains an ubiquitin C promoter (UBC) promoter that drives the constitutive expression of the reverse tetracycline transactivator 3 (rtTA3) protein as well as a puromycin (Puro) selectable marker located 3’ to an internal ribosome entry site (IRES). (C) Mature BHRF1 miRNA expression was analyzed by miRNA-specific Taqman qPCR. Expression levels in ∆123 LCLs transduced with the pTREX–based miR-BHRF1-123 expression vector was normalized to the matched WT LCL donor. Average of two donors with SD indicated. No miR-BHRF1 miRNAs were detected in ∆123 LCLs transduced with the parental pTREX vector. (D) LCLs were transduced with either the empty pTREX vector or the miR-BHRF1-123 expressing pTREX vector. After puro selection, total Thy1.1+ cells were counted every 3 days and normalized to WT cell total. Average of two donors. Significance determined by two-way ANOVA with multiple comparisons. *p<0.05. Error bars=SD.

We next wanted to establish that the miR-BHRF1 miRNAs were indeed necessary and sufficient to rescue the slow growth phenotype of ∆123 LCLs by ectopic expression of the entire repertoire of miR-BHRF1 miRNAs. To this end, the miR-BHRF1-1, 1–2 and 1–3 pri-miRNAs were cloned into the antisense-oriented intron present in the Tet-inducible lentiviral miRNA expression vector pTREX, as this vector allows a consistently higher level of pri-miRNA expression when compared to conventional lentiviral vectors (Fig.1B) (Poling et al., 2017). miR-BHRF1-1 and 1–3 are not well expressed from their natural pri-miRNA stem-loops, isolated from the EBV genome (Feederle et al., 2011a; Haar et al., 2015), so they were expressed from expression cassettes based on the human pri-miR-30 precursor, as previously described (Zeng et al., 2002). Additionally, miR-BHRF1-2/2* was cloned into pTREX using its natural stem-loop and flanking sequences. After selection of transduced cells by puromycin selection, and induction of miRNA expression by addition of doxycycline (Dox), we observed levels of expression of miR-BHRF1-1, 1–2, 1–2* and 1–3 in ∆123 LCLs that were similar to the expression levels observed in WT LCLs (Fig. 1C). Since miRNAs are thought to regulate mRNA expression in *trans* (Chong et al., 2010; Guo et al., 2010; Zeng et al., 2002), we hypothesized that ectopic expression of the miR-BHRF1 miRNAs in ∆123 LCLs would increase the growth of these mutant LCLs to WT levels. However, we did not observe any rescue of the slow growth phenotype (p<0.05) (Fig. 1D). This suggested that the miR-BHRF1 miRNAs do not enhance LCL growth via their canonical role of mRNA downregulation in *trans* by translational inhibition or degradation of targeted mRNAs (Chendrimada et al., 2005; Huntzinger and Izaurralde, 2011; Liu et al., 2004; Meister et al., 2004).

### The EBNA-LP protein and co-transcriptional activity are upregulated in ∆123 LCLs

Because the slower growth of the ∆123 LCLs was not rescued by expression of the EBV miR-BHRF1 miRNAs in *trans*, we sought an alternative hypothesis to explain this phenotype. Previously, we reported that ∆123 LCLs display a marked upregulation in the expression of the EBNA-LP protein (Feederle et al., 2011a, 2011b) and we confirmed that EBNA-LP levels in ∆123 LCLs are indeed between 5- and 20-fold higher than seen in control WT LCLs (Fig. 2). This latter result cannot be explained based on the model that the miR-BHRF1 miRNAs are acting to downregulate EBNA-LP mRNA expression in *trans,* as there are no predicted target sites for these viral miRNAs in the EBNA-LP mRNAs, and moreover we and others failed to detect the binding of miR-BHRF1 miRNAs to EBNA-LP mRNAs, or the viral HF exon, using high throughput CLIP techniques (Riley et al., 2012; Skalsky et al., 2012).

**Figure 2.**
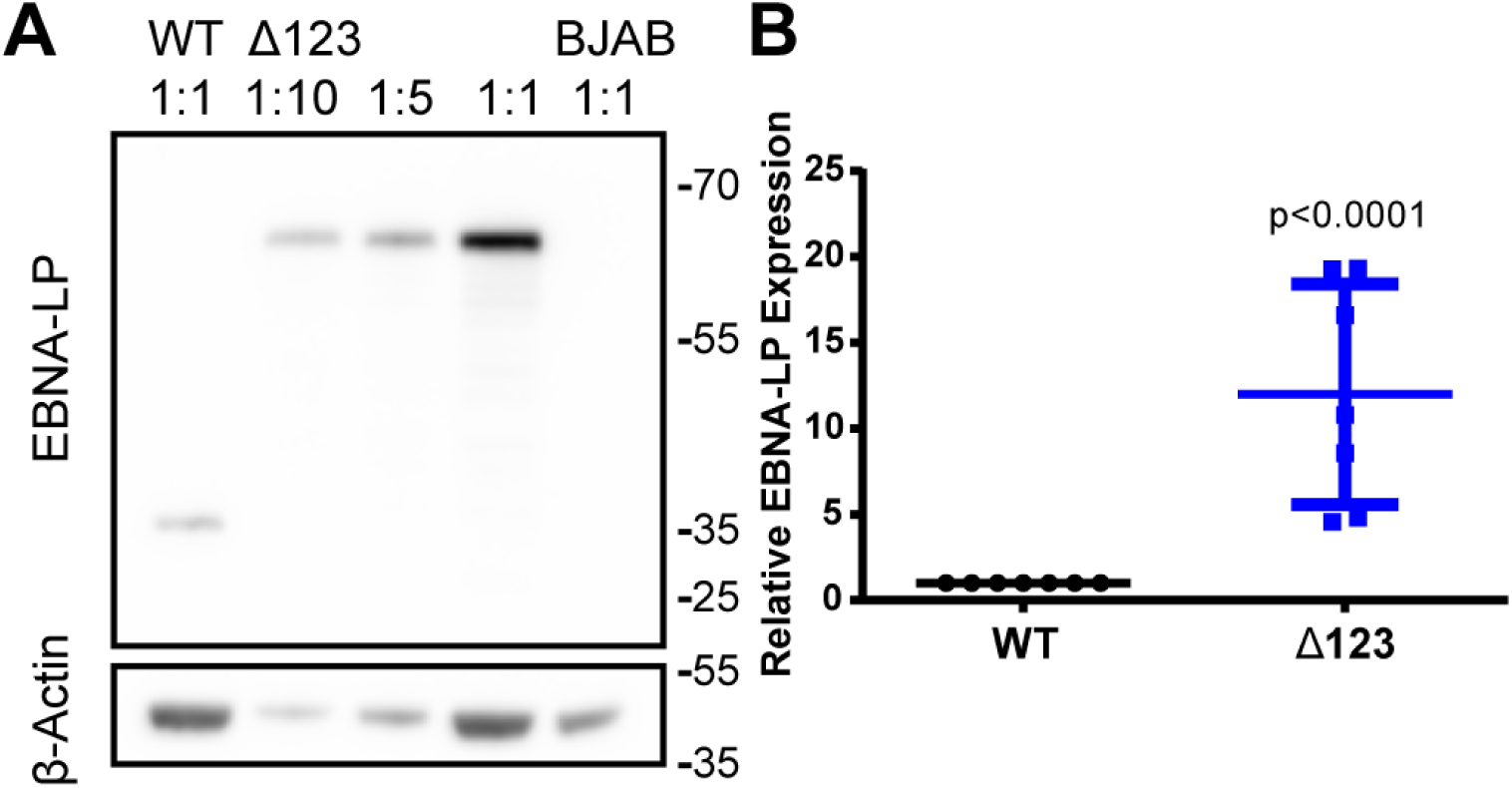
EBNA-LP is significantly overexpressed in ∆123 LCLs. (A) Western blot of EBNA-LP expression in WT and ∆123 LCLs. The ∆123 LCL lysate was loaded at increasing concentration left to right, as indicated. BJAB lysate was loaded as a negative control. β-Actin was used as a loading control. (B) ∆123 LCL EBNA-LP expression normalized to WT LCL levels. Average of seven donors. p < 0.0001; one-sample t-test. Error bars = SD.

To determine if the increased expression of the EBNA-LP protein correlated with an upregulation of the Cp and LMP1 promoters, which are transcriptionally activated by EBNA-LP and EBNA2 (Harada and Kieff, 1997; Nitsche et al., 1997; Peng et al., 2005; Sung et al., 1991; Wang et al., 1990; Woisetschlaeger et al., 1990), we used 4-thiouridine (4SU) to label nascent transcripts for 60 min. Nascent transcripts were purified and their expression levels were then compared to total transcript levels to determine half-life, nascent transcript expression, and total transcript expression (Fig. 3A-C). As expected, we saw a significant increase in the level of nascent and total Cp driven and LMP1 transcripts in the ∆123 LCLs, when compared to control mRNAs encoding GAPDH, β-Actin, and SETDB1 (Fig. 3A and B), indicating that overexpression of EBNA-LP leads to a specific increase in EBNA-LP-mediated transcriptional activation of these viral mRNAs. In contrast, we did not see any major change in viral mRNA half-life when WT LCLs were compared to ∆123 LCLs (Fig. 3C). In addition to the upregulation of EBNA-LP regulated transcripts, we also saw a significant increase in both total and nascent Wp driven transcripts for some of our donors, which corroborates published findings (Feederle et al., 2011a, 2011b).

**Figure 3.**
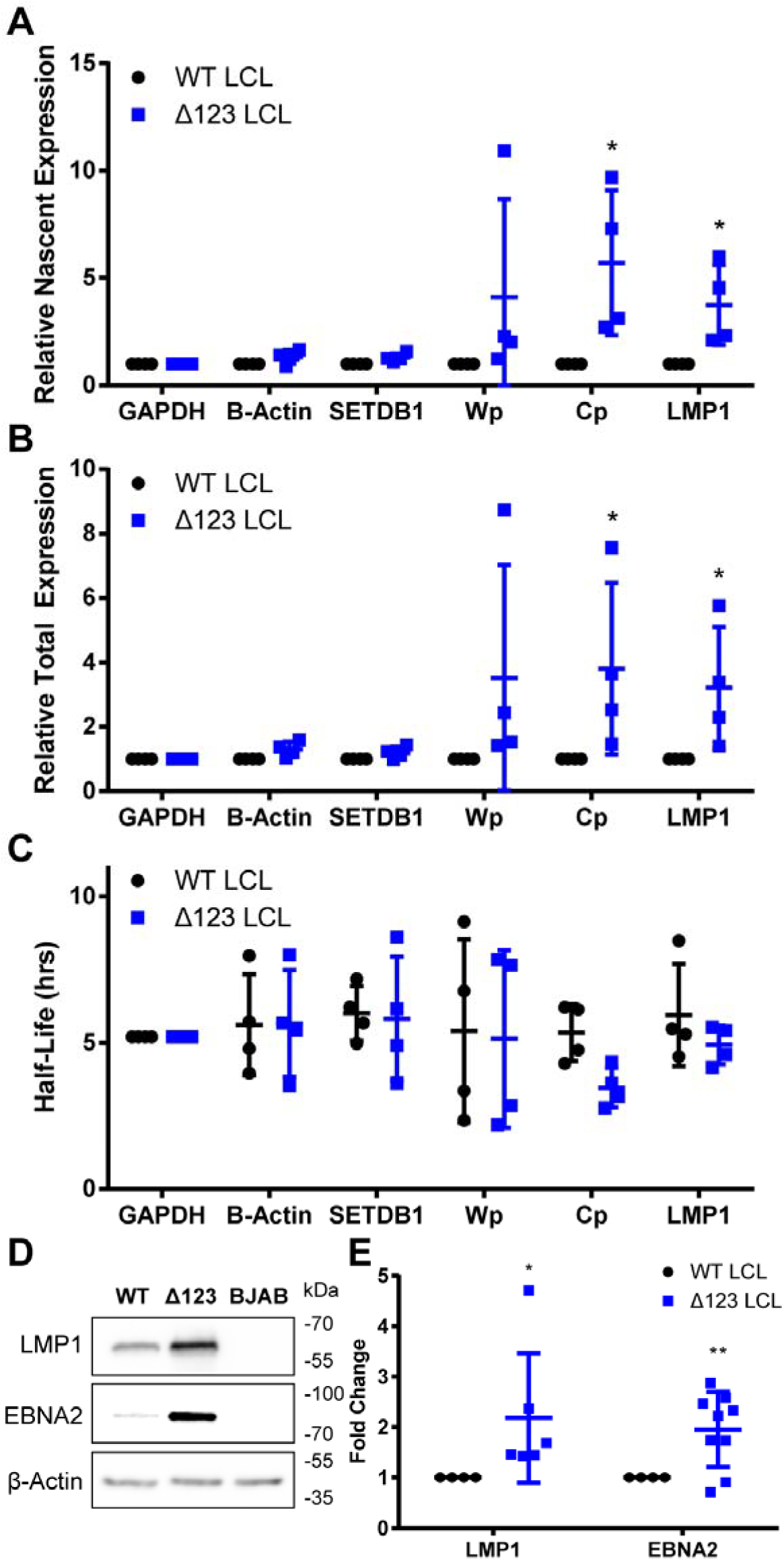
∆123 LCLs show an increase in EBNA-LP regulated transcription and protein expression. Transcript expression was tested by treating LCLs with 200 uM 4SU for 1 h. Total RNA was harvested and 4SU incorporated into nascent RNA labeled with biotin and isolated using magnetic streptavidin beads. Nascent and total RNA levels were measured by qPCR and used to calculate (A) relative nascent transcription, (B) relative total transcription, and (C) half-life. GAPDH was used as a normalization control and β-Actin and SETDB1 were used as negative controls. Wp, W promoter driven transcripts. Cp, C promoter driven transcripts, LMP1, and CCND2 are known to be co-transcriptionally activated by EBNA-LP. *p<0.05, one-sample t-test. Error bars = SD. (D) Western blot of EBNA-LP-regulated viral proteins LMP1 and EBNA2 from WT and ∆123 LCLs. BJAB lysate was used as a negative control for viral proteins. (E) Quantitation of Western blot analysis normalized to WT expression levels. Data represent the average of 6+ donors. *p<0.05, ** p<0.01; one-sample t-test. Error bars = SD.

In addition to an increase in EBNA-LP regulated viral transcripts, the encoded viral proteins were also upregulated in ∆123 LCLs. Thus, we saw an upregulation of LMP1 expression in all six donors tested and upregulation of EBNA2 in six out of eight donors tested (Fig. 3D and 3E). The EBNA2 exon (YH) is one of many exons that can be alternatively spliced in Cp and Wp-driven pre-mRNAs; however, during the establishment of latency, the Cp promoter, due to activation from EBNA2 and EBNA-LP, becomes the dominant source of EBV transcription (Harada and Kieff, 1997; Sung et al., 1991; Woisetschlaeger et al., 1990). The two donors without an upregulation of EBNA2 may alternatively splice to non-YH containing exons at an increased level, and not to YH itself, leading to a basal expression level of EBNA2. Nevertheless, EBNA2 is upregulated in the ∆123 LCLs derived from 75% of the donors tested (Fig. 3E), which is statistically significant (p<0.01).

### The miR-BHRF1 microRNAs downregulate EBNA-LP expression in *cis*

We next sought to determine how disruption of miR-BHRF1 miRNA expression induces overexpression of EBNA-LP. Early reports indicated that the HF exon, which encompasses the miR-BHRF1-2 and miR-BHRF1-3 pri-miRNAs, forms the 3’UTR of a minor portion of EBNA-LP mRNAs (Austin et al., 1988; Bodescot and Perricaudet, 1986; Pfitzner et al., 1987). However, the question of whether these pri-miRNAs affect EBNA-LP mRNA expression in *cis* has not been previously addressed. We therefore sought to determine if the BHRF1 miRNAs are indeed encoded within the 3’UTR of EBNA-LP mRNAs, and if so, how often this occurred. In this context, we were particularly interested in a previous report (Lin and Sullivan, 2011) demonstrating that the human γ-herpes KSHV uses viral pri-miRNAs to downregulate expression of the viral Kaposin B mRNA and protein in *cis*.

The miR-BHRF1 miRNAs are encoded within two exons. The more 5’ exon (H2) contains the miR-BHRF1-1 pri-miRNA while the more 3’ exon (HF) contains the BHRF1 protein coding sequence and the miR-BHRF1-2 and 1–3 pri-miRNAs in the BHRF mRNA 3’UTR (Fig. 4A). To determine if the H2 or HF exons form part of the 3’UTR of EBNA-LP transcripts, we performed a meta-analysis of RNA-Seq data from 10 random donors from the 1000 genomes project (Fig. 4B) (Lappalainen et al., 2013). Reads that failed to align to the human genome were aligned to the WT EBV genome (Accession: NC_007605) using the hierarchical indexing for spliced alignment of transcripts (HISAT2) splice-aware RNA-Seq alignment software (Kim et al., 2016). Splice junctions were identified and quantified by visualizing data in the integrative genome viewer (IGV) (Robinson et al., 2011). These alignments revealed no splice junctions from any Cp or Wp-driven exon located 5’ of H2 to the H2 exon. This suggests that miR-BHRF1-1 is not encoded within any of the mature transcripts driven from the Cp and Wp promoters and thus cannot form part of the EBNA-LP mRNA 3’UTR. Instead, and as previously proposed, miR-BHRF1-1 is likely excised from an intronic location (Cai et al., 2006; Xing and Kieff, 2007). In contrast, our data do provide strong evidence in support of the hypothesis that both the Y2 exon and, to a lesser extent the Y3 exon, is frequently spliced to the HF exon (Fig. 4B). In particular, Y2-HF splice junction read depth was comparable to those of canonical EBV splice junctions, such as Y2 to U and Y3 to U (Fig. 4B). These data suggest that the BHRF1 ORF, and the miR-BHRF1-2 and 1–3 pri-miRNA stem-loops, likely constitute a high percentage of the 3’UTRs of EBNA-LP mRNAs. However, because of the differential use of the alternative splice acceptors W1 and W1’, where W1’ is necessary to generate the ATG translation initiation codon for EBNA-LP expression, it remained unclear if any functional EBNA-LP mRNAs indeed contain the EBNA-LP ORF located 5’ to the HF exon (Fig. 4A).

**Figure 4.**
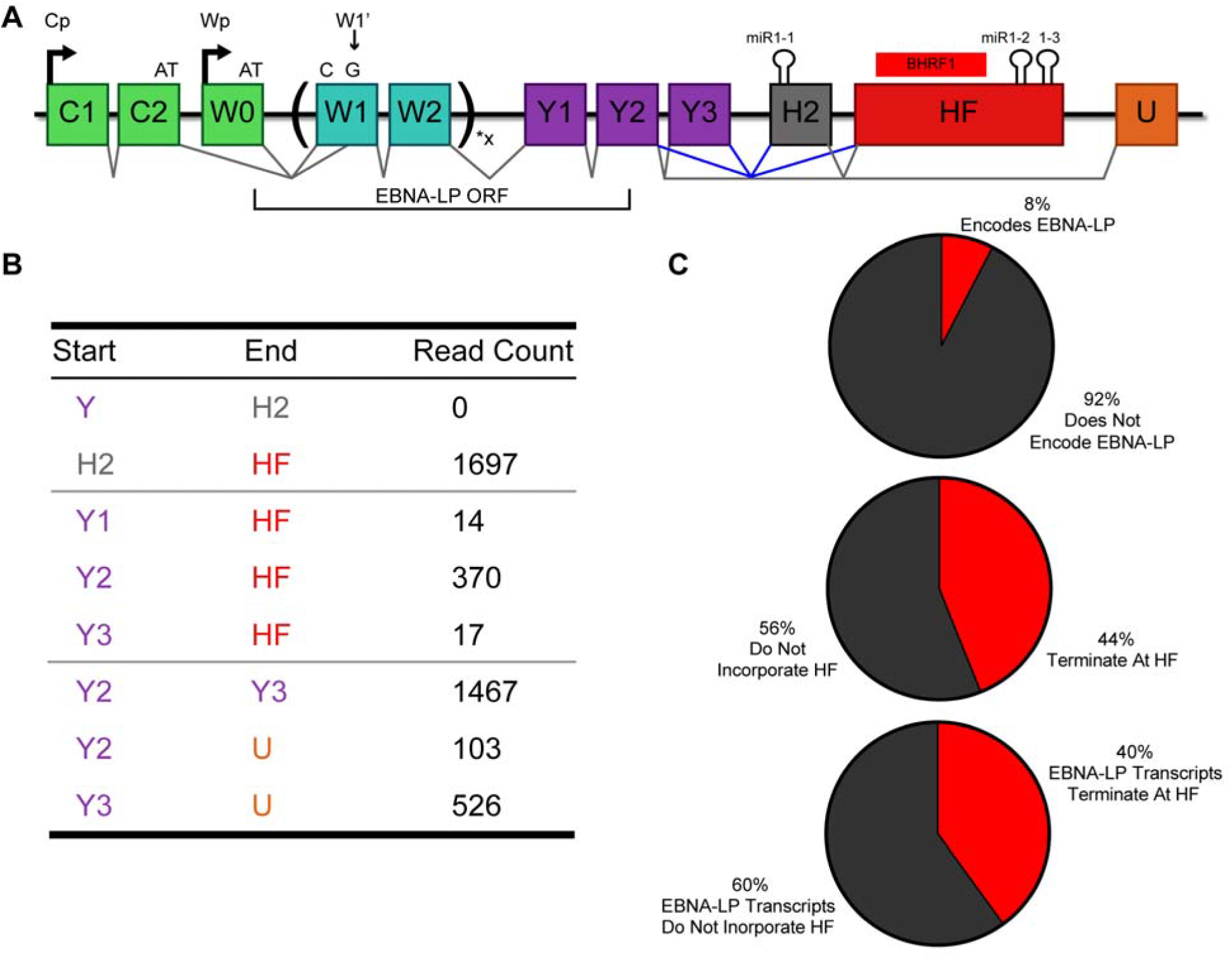
EBNA-LP encoding transcripts are alternatively spliced to include the HF exon at a high frequency. (A) Schematic of EBNA-LP alternative splicing. EBNA-LP mRNA transcription is driven by either the Cp or Wp promoter. Both promoters encode an exon that terminates in AT (C2 or W0) that must splice to a 3’ splice site located within the first W1 exon, called W1’, to acquire the G to make the ATG necessary to initiate EBNA-LP translation. EBNA-LP transcripts then splices to W2 and can include anywhere from 1–11 copies of the W1/W2 repeats. The coding region for EBNA-LP then terminates within the Y2 exon. The H2 exon contains the BHRF1-1 pri-miRNA and the HF exon contains the BHRF1 ORF and pri-miR-BHRF1-2 and 1–3. Known splice junctions are shown in black, including to the U exon that splices 3’ to exons encoding EBNA1, 3A, 3B and 3C. Possible splice junctions that would include the BHRF1 miRNAs in the 3’UTR of EBNA-LP are shown in blue. (B) RNA-Seq data from the 1000 genomes project was processed to identify splice junctions. Raw read counts are shown for H2 and HF splicing. Canonical splice junctions are provided at the bottom for reference. (C) PacBio sequencing was performed on RNA isolated using a probe complementary to the W2 exon. Of the Cp and Wp driven transcripts sequenced, the percentage that encode EBNA-LP, that terminate with the HF exon, and EBNA-LP mRNAs that terminate with the HF exon, are shown.

Since EBNA-LP-encoding transcripts that terminate at the HF exon can only be identified in long sequencing reads that encompass not only the HF exon but also the 5’ end of the EBNALP ORF, we performed PacBio sequencing for EBV transcripts expressed in WT EBV strain B95.8-derived LCLs. Total RNA was harvested and EBV specific transcripts were enriched using a biotinylated antisense oligonucleotide specific for the W2 exon common to all EBNA transcripts followed by pull-down using streptavidin beads (Fig. 4A). This enriched pool of EBV transcripts was then oligo-dT primed to generate long cDNAs that were sequenced using a PacBio RSII analyzer. We were able to enrich the Wp/Cp driven EBV transcripts by ~7,000-fold over PacBio sequencing previously published from total LCL mRNA (O’Grady et al., 2016; Tilgner et al., 2014). EBV transcripts make up a low percentage of total LCL transcripts, as seen in our RNA-Seq meta-analysis (EBV transcripts were only ~0.70% of total reads). Therefore, despite our 7,000-fold enrichment, we identified 120 reads that mapped to EBV, 95 reads that mapped to the Wp/Cp transcript region, 29 reads that contain the HF exon, and 5 reads that encode EBNA-LP (Fig. 4C). However, these data do indicate that the HF exon is present in a high proportion of all Wp/Cp-driven transcripts (44%), and, more importantly, in EBNA-LP-encoding transcripts (40%). This indicates that excision of the pri-miR-BHRF1 miRNAs located in the HF exon has the potential to significantly reduce EBNA-LP transcript levels, and hence EBNA-LP expression.

Cleavage of a pri-miRNA stem-loop by the microprocessor complex, consisting of Drosha and DGCR8, results in excision of the pre-miRNA and degradation of the 5’ and 3’ pre-miRNA flanking regions (Cai et al., 2004). Therefore, the excision of pre-miR-BHRF1-2 and 1–3 is predicted to reduce EBNA-LP protein expression. To test this hypothesis, we cloned the 3’UTR of the BHRF1 protein, which encompasses these two pri-miRNAs, either from WT EBV B95-8 or from the ∆123 EBV mutant in which the pri-miR-BHRF1-2 and 1–3 stem-loops have been mutationally inactivated, 3’ to the FLuc indicator gene. These two constructs, and a control “empty” FLuc vector lacking any inserted EBV sequences, were then transfected into wildtype 293T cells, as well as into a previously described 293T derivative in which both Drosha and Dicer expression had been blocked by gene editing, called RNaseIII-/- cells (Aguado et al., 2017). FLuc expression was then determined relative to a co-transfected NanoLuc control vector. The ratio of the FLuc expression seen with the empty FLuc vector in WT 293T cells versus the RNaseIII-/-cells was then set at 1.0. As shown in Fig. 5A, we see a significant ~4-fold reduction of FLuc activity in the WT 293T cells, relative to the RNaseIII-/- cells, when pri-miR-BHRF1-2 and 1–3 were present in the 3’UTR, and this inhibition was fully rescued when the two pri-miRNAs stem-loops were mutated (p<0.01). We also confirmed the efficient excision of pre-miR-BHRF1-2 and 1–3 from the FLuc-based reporter construct by visualizing the expression of the resulting miRBHRF1-2, 1–2* and 1–3 miRNAs by Northern blot analysis (Fig. 5B).

**Figure 5.**
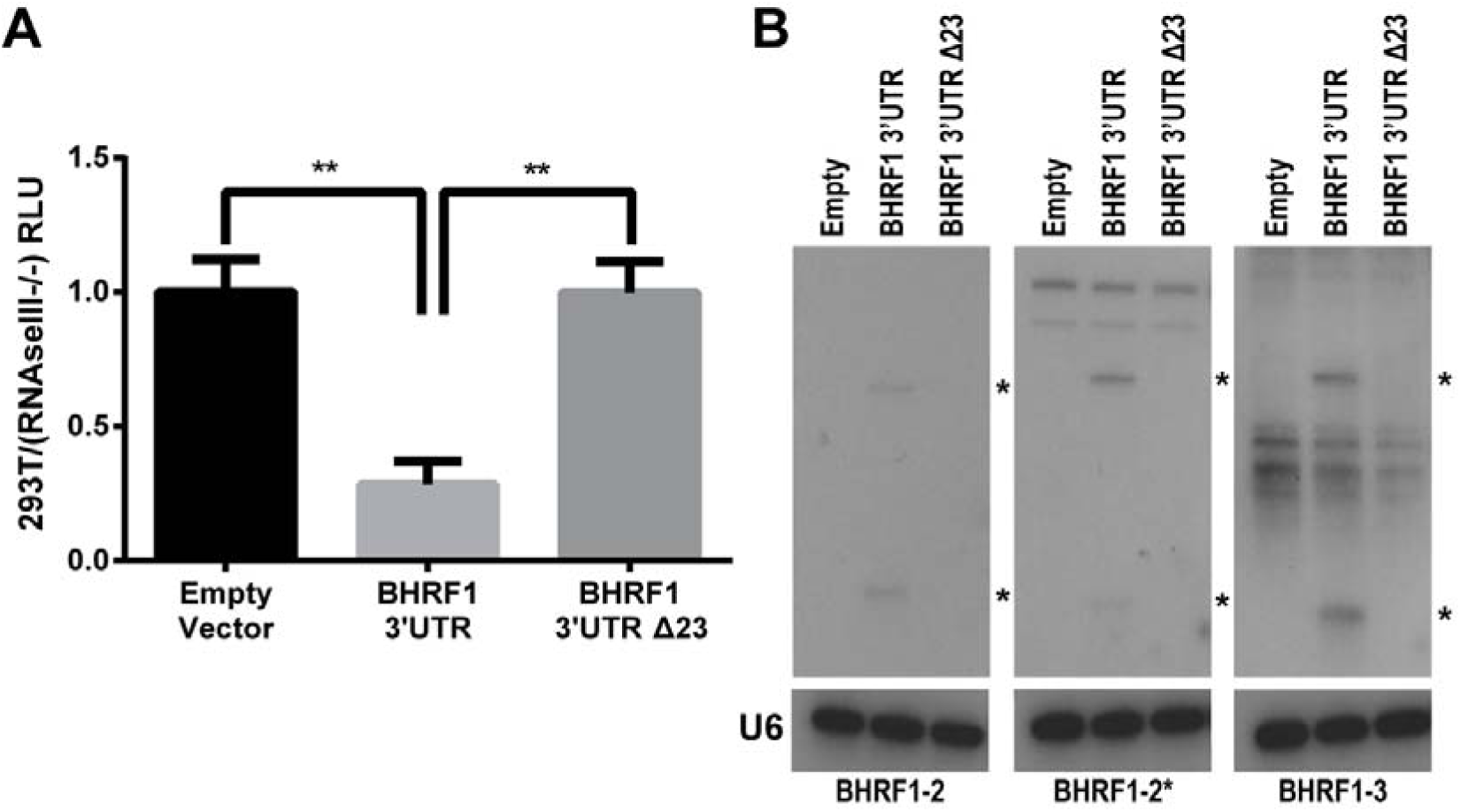
pre-miR-BHRF1-2/2* and 1–3 miRNA excision regulates protein expression in *cis*. (A) The BHRF1 3’UTR, containing the pri-miR-BHRF1-2 and 1–3 stem-loops (Fig. 4), was cloned from both the WT and ∆123 EBV virus into the 3’UTR of a FLuc expression vector. These vectors, along with a NanoLuc internal control vector, were transfected into both 293T cells and RNaseIII-/- cells and luciferase activity measured. Expression level was normalized to NanoLuc followed by an empty FLuc vector control. Relative 293T luciferase expression was normalized to the expression detected in the RNaseIII-/- cells to observe differences due to Drosha processing. Five biological replicates were performed. **p<0.01, one-sample t-test, and student’s t-test. Error bars = SD. (B) Northern blots show miRNA expression from the BHRF1 3’UTR firefly luciferase vector. Blots were probed with a U6-specific probe to demonstrate equal loading. * Indicates pre- and mature miRNAs.

To further confirm the presence of the HF exon in a least a subset of EBNA-LP mRNAs, we generated 12 artificial miRNAs (amiRNAs) that were designed to target the HF exon and identified two amiRNAs, called B4 and B11, that cleaved the HF exon effectively. ∆123 LCLs were then transduced with a pTREX-based lentivector expressing either B4 or B11 and a Western blot analysis for EBNA-LP expression performed on puromycin selected, Dox-induced cells. We indeed saw a reduction of EBNA-LP protein expression when either HF targeting amiRNAs was expressed (Fig. 6). The observed reduction in EBNA-LP protein when the HF exon was targeted by RNAi confirms that at least some EBNA-LP mRNAs contain HF and suggest that destabilization of these EBNA-LP transcripts by excision of the pri-miR-BHRF1-2 and 1–3 is indeed likely to occur.

**Figure 6.**
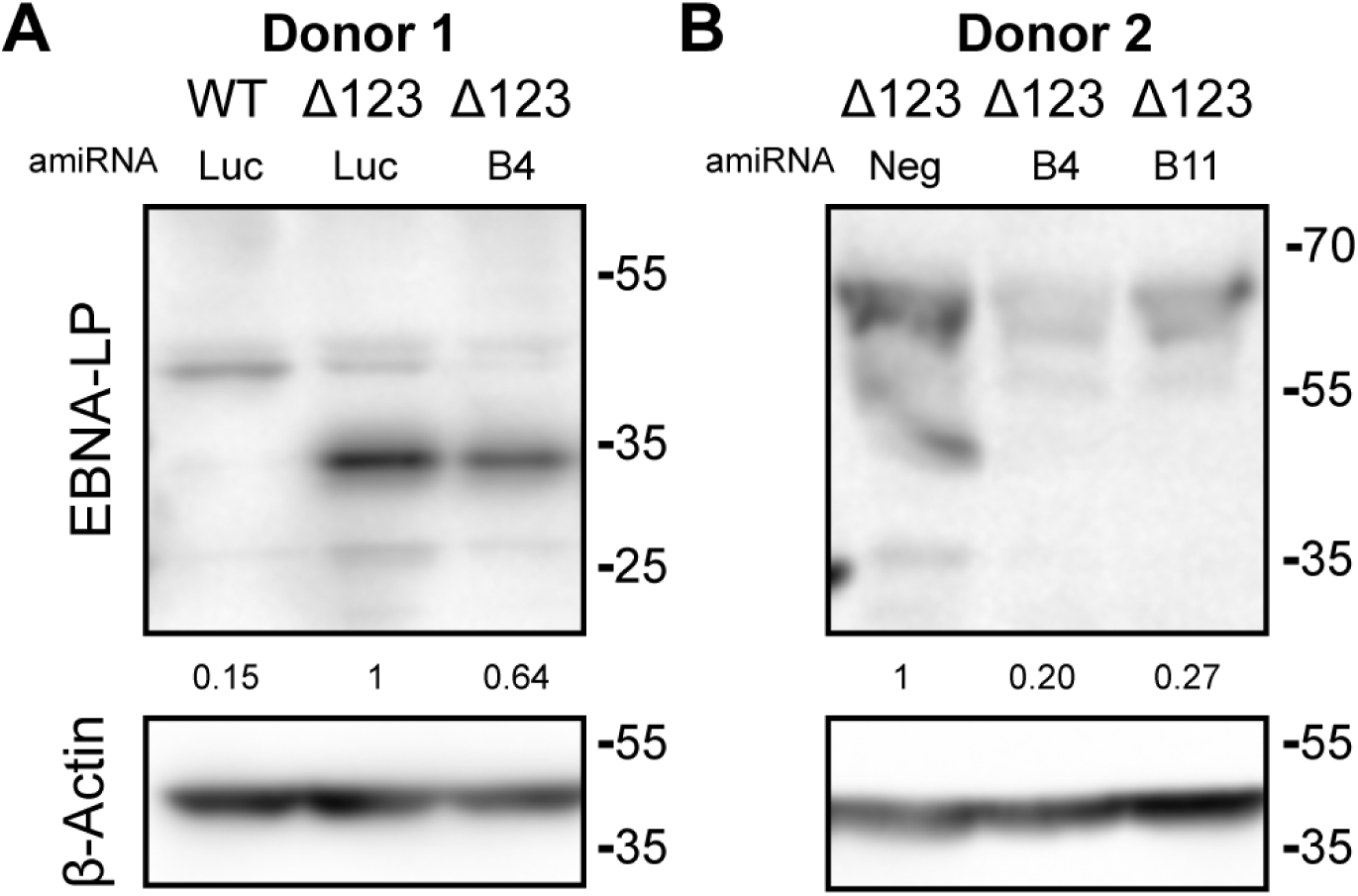
amiRNAs targeting the HF exon reduce EBNA-LP expression. (A) Donor one: WT and ∆123 LCLs were transduced with a control FLuc (luc) amiRNA. The ∆123 LCLs were also transduced with an amiRNA targeting the HF exon (B4). (B) Donor 2: only the ∆123 LCLs were transduced with two amiRNAs targeting the HF exon (B4 and B11). These transduced LCLs are compared to the originating, non-transduced ∆123 LCLs. Transduced cells were selected with puro and then induced with Dox for 4 days before lysates were harvested. Western blot analysis was then performed for EBNA-LP. β-Actin was used as a loading control. Fold knockdown relative to ∆123 LCL expression is shown for EBNA-LP.

### PacBio sequencing reveals additional novel Wp/Cp splice patterns

PacBio sequencing is a powerful tool for elucidating novel transcription patterns due to its ability to sequence full-length mRNAs. By performing PacBio sequencing on EBV transcripts, we identified two novel EBV RNA splicing patterns. First, we observed a new 3’ splice site within the C2 exon, which we term C2’, that was observed in 35% of sequenced transcripts containing C2 (Fig. 7A and 7B). This splice donor, when used, excludes the final seven base pairs of the C2 exon. This is especially relevant to the EBNA-LP ORF, since it eliminates the AT dinucleotide necessary for EBNA-LP translation initiation. Since the C2’ 3’ splice site was used 35% of the time, this novel splice junction contributes to the low level of Cp/Wp transcripts capable of encoding EBNA-LP (8%) (Fig. 4C and Fig. 7B). We looked for the C2’ splice junction in our meta-analysis of the 1000 genomes RNA-Seq data (Lappalainen et al., 2013) and observed that the C2’ splice site is used 27% of the time, validating our PacBio sequencing data. We also observed splicing from the 5’ splice site at the end of the W2 exon to a W1’ 3’ splice site. This W2-W1’ splicing event occurs five base pairs internal to W1, resulting in a frame shift in the EBNA-LP ORF and causing a premature stop codon to occur, thereby inhibiting the expression of EBNA-LP. Since this internal W1’ splicing event occurs 48% of the time, it would cause significantly reduced EBNA-LP expression. Combined, these two novel splicing events prevent the expression of EBNA-LP by a significant number of Cp/Wp initiated transcripts.

**Figure 7.**
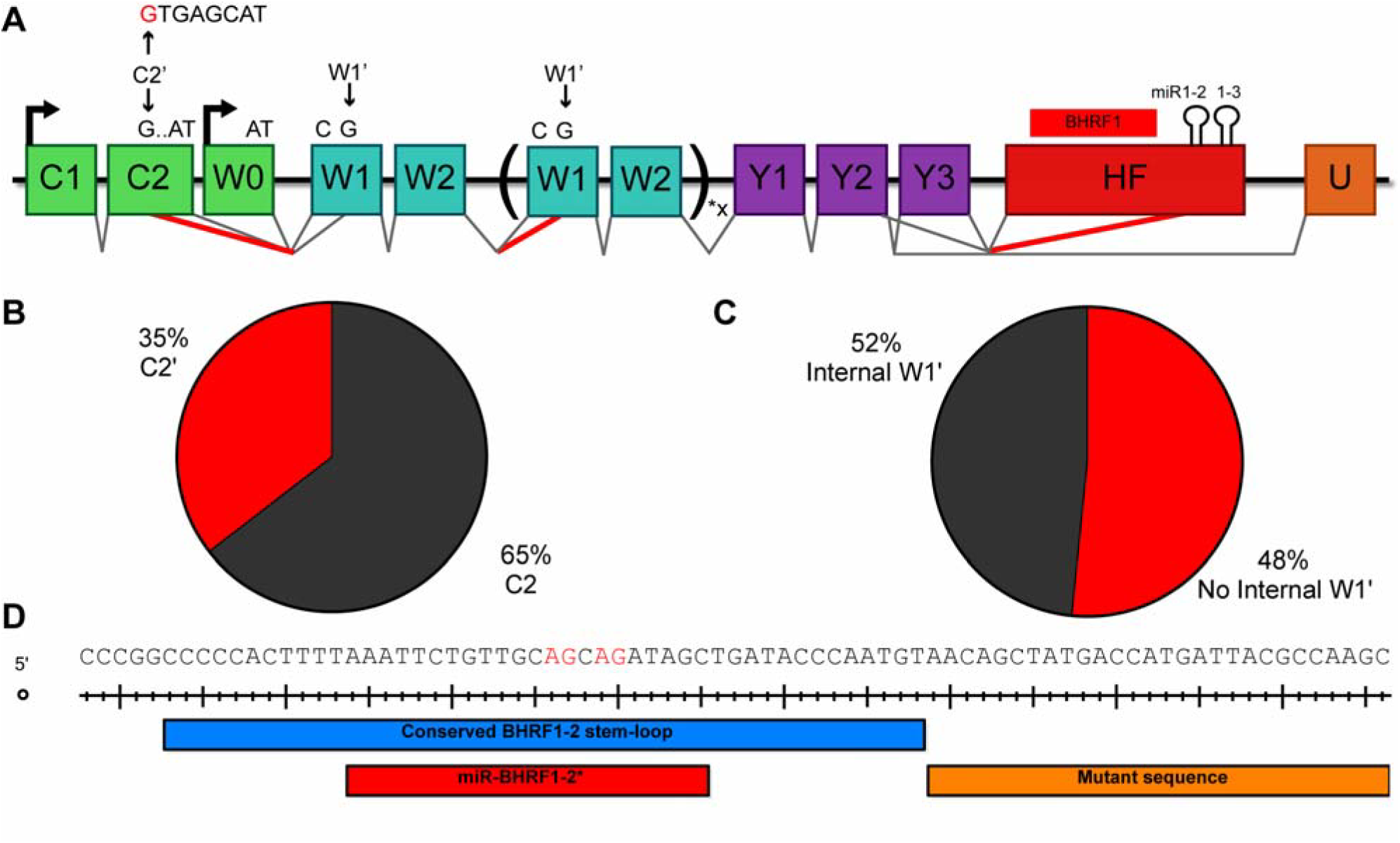
Novel alternative splicing patterns of Cp and Wp transcripts. (A) Schematic of all novel splice patterns observed in EBV strain B95-8 LCLs by PacBio sequencing. There is a novel 5’ splice site in C2, termed C2’, that is shown in red. Use of this splice site removes the last seven bases of C2. Splicing was also observed between W2 and W1’, generating a frame shift in the EBNA-LP ORF that causes premature translation termination. The ∆123 LCLs exhibit cryptic 3’ splice sites in the mature miR-BHRF1-2 sequence, shown in red. This results in efficient splicing from the Y2 and Y3 exons to the miR-BHRF1-2 stem-loop. Percentages exhibited in the PacBio sequencing are shown for (B) C2’ usage, (C) internal W2-W1’ splicing. (D) Cryptic splice acceptors sites are shown in red. The portion of the pri-miR-BHRF1-2 stem-loop retained in the ∆123 EBV mutant is indicated in blue, the mature miR-BHRF1-2* sequence is shown in red, and the mutant sequence introduced by the ∆123 deletion of the 3’ side of the BHRF1-2 pri-miRNA is shown in orange.

The third novel EBV splicing pattern we observed in this study resulted from shallow RNA sequencing of the ∆123 LCLs. With the goal of establishing that EBNA-LP coding transcripts terminate in the HF exon, we performed shallow sequencing on both WT and ∆123 LCLs. In doing so, we noticed splicing from the Y2 exon to two closely adjacent 3’ splice sites internal to the HF exon, specifically within the mature miR-BHRF1-2* sequence. The deletion mutation introduced into the ∆123 EBV mutant removed the 3’ arm of the pri-miR-BHRF2 stem-loop while leaving the 5’ arm, including the mature miR-BHRF2* sequence, intact. Even though the miR-BHRF1-2* sequence is, of course, present in WT EBV strain B95-8, we did not detect this splicing event in either our shallow sequencing or PacBio sequencing of WT LCL transcripts. We hypothesize that this is because these two novel 3’ splice sites are not normally available for splicing due to their sequestration within the pri-miR-BHRF1-2 stem-loop in LCLs generated using WT EBV. Whether this novel splicing event has any effect on EBV gene expression during either the latent or lytic phase of the viral replication cycle is unclear, though it clearly has the potential to inhibit BHRF1 protein expression.

## Discussion

Previous studies by our laboratory and others demonstrated that the elimination of the miR-BHRF1 miRNAs in the ∆123 EBV mutant inhibited B cell transformation and reduced the growth rate of the subsequent ∆123 LCL cell lines (Feederle et al., 2011a, 2011b; Haar et al., 2015; Seto et al., 2010). In this study, we show that ectopic expression of physiological levels of the miR-BHRF1 miRNAs does not restore rapid growth to ∆123 LCLs (Fig. 1C). To explain this result, we propose a model whereby excision of pre-miR-BHRF1-2 and 1–3 from the EBNA-LP mRNA 3’UTR by the microprocessor complex is important for expression of the appropriate level of EBNA-LP, and hence of EBNA-LP regulated proteins (Fig. 8).

**Figure 8.**
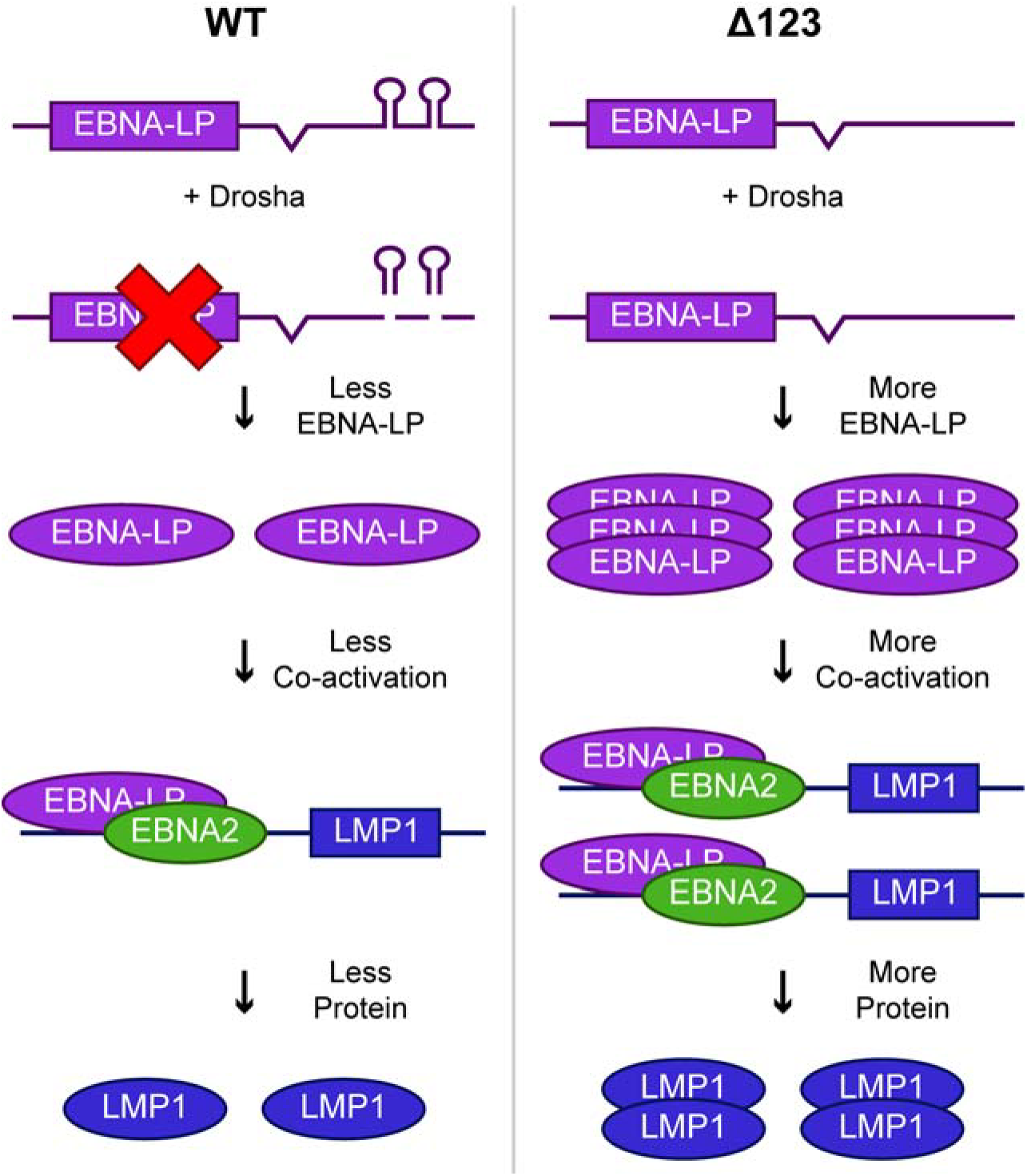
Model for how the pri-miR-BHRF1-2 and 1–3 miRNAs regulate EBV gene expression in *cis*. In the context of WT EBV infection (left side), pri-miR-BHRF1-2 and 1–3 form part of the 3’UTR of EBNA-LP mRNAs. Cleavage of these pri-miRNAs by Drosha leads to a decrease in EBNA-LP transcript levels and, hence, EBNA-LP transcription factor expression. This in turn leads to a lower level of expression of EBNA-LP regulated transcripts, such as LMP1, and their encoded proteins. In contrast, when pri-miR-BHRF1-2 and 1–3 are deleted (right side), in the EBV ∆123 mutant, EBNA-LP and its downstream viral gene targets are overexpressed, resulting in reduced B cell transformation and slower LCL growth.

While it is clear that the BHRF1 miRNAs play a role in B cell transformation and growth, their mechanism of action has remained unclear. However, given that the canonical mechanism of action of miRNAs is to repress mRNA function in *trans*, and given that mRNA targets bound by the miR-BHRF1 miRNAs can be readily identified by high throughput CLIP approaches (Riley et al., 2012; Skalsky et al., 2012), it seemed likely that these miRNAs were acting to repress the expression of cellular factors that inhibit B cell transformation. By studying the individual roles of BHRF1 miRNAs in growth and transformation, previous research has revealed that a mutant EBV lacking only miR-BHRF1-1 is fully competent for LCL transformation (Feederle et al., 2011a). Moreover, mutation of the BHRF1-2 and 1–2* seed sequences, while leaving the structure of the pri-miR-BHRF2 stem-loop intact, also revealed no difference in B cell transformation efficiency, and LCL cell cycle progression, despite the fact that a mutant EBV virus in which the pri-miRBHRF1-2 stem-loop was deleted demonstrated markedly reduced transformation efficiency (Feederle et al., 2011a; Haar et al., 2015). The Delecluse laboratory has also shown that pri-miRBHRF1-2 processing is necessary for efficient processing and expression of pri-miR-BHRF1-3 and they therefore proposed, essentially by a process of elimination, that loss of miR-BHRF1-3 expression is solely responsible for the decrease in B cell transformation and LCL growth seen with the ∆123 EBV mutant (Haar et al., 2015). This was surprising as the seed sequence of miRBHRF1-3, unlike the seed sequence of miR-BHRF1-2, is not conserved in primate lymphocryptovirus (LCV) evolution (Cai et al., 2006; Riley et al., 2010; Skalsky et al., 2014). However, we show here that ectopic expression of all of the miR-BHRF1 miRNAs, including miRBHRF1-3, in ∆123 LCLs fails to restore the rapid growth seen with WT LCLs (Fig. 1D). The apparent requirement for the pri-miR-BHRF1 stem-loops, but not their activity as miRNAs, for optimal LCL growth suggested the alternative hypothesis that the EBV pri-miR-BHRF1 stem-loops promote B cell transformation and LCL growth by acting in *cis* to promote the expression of an optimal level of the EBNA-LP transcription factor and, hence, of its downstream gene targets.

As previously reported, we indeed observe a marked increase in EBNA-LP protein expression in ∆123 LCLs, relative to WT LCLs (Fig. 2) (Feederle et al., 2011a, 2011b). As would be predicted, we also saw an increase in the expression of EBNA-LP regulated viral transcripts and proteins, specifically Cp and LMP1p-driven transcripts and the EBNA2 and LMP1 proteins (Fig. 3). CLIP analysis has revealed no target sites for the miR-BHRF1 miRNAs in Cp or Wp-driven transcripts, suggesting that the miR-BHRF1 miRNAs do not directly inhibit Cp/Wp transcript expression (Riley et al., 2012; Skalsky et al., 2012). Early reports identified rare Cp/Wp transcripts with splicing from the Y exons to the HF exon (Austin et al., 1988; Bodescot and Perricaudet, 1986; Pfitzner et al., 1987) and we therefore hypothesized that the miR-BHRF1 pri-miRNAs might be located in the EBNA-LP 3’UTR. This would be expected to result in an inhibition of EBNA-LP mRNA expression in *cis,* due to destabilization resulting from the excision of the pre-miRNAs by microprocessor cleavage, and thus provided an attractive alternative explanation for the apparent inverse correlation between EBNA-LP and miR-BHRF1 miRNA expression. Previous research provides evidence that splicing of Cp/Wp transcripts to the HF exon does indeed occur. Thus, Arvey et. al. showed by RNA-Seq that splicing from the Y2 exon to HF occurred in 35% of all Cp/Wp driven transcripts (Arvey et al., 2012), while Tierney et al. showed by qRT-PCR that Y2-HF transcripts are highly expressed in LCLs (Tierney et al., 2015). However, due to the technical limitations of RNA-Seq and qRT-PCR, these studies could not confirm that the viral sequences located 5’ to the HF exon indeed encoded EBNA-LP rather than a long 5’UTR on a BHRF1 mRNA, for example. Our study expands on this previous work by looking at long EBV sequence reads generated by PacBio sequencing. From this, we determined that ~44% of our Cp/Wp transcripts contain the HF exon as their 3’UTR and ~40% of transcripts bearing an intact EBNA-LP ORF terminate within the HF exon (Fig. 4C). Previous PacBio sequencing of LCLs did not include any enrichment for Cp/Wp-driven transcripts, and so failed to detect any splicing between the Y exons and the HF exon in latent transcripts (O’Grady et al., 2016). We have therefore identified a novel, yet common, splicing pattern for EBNA-LP transcripts that allows the miR-BHRF1-2 and 1–3 pri-miRNA stem-loops to downregulate EBNA-LP expression in *cis*. Importantly, we note that excision of miR-BHRF1-2 and/or 1–3 from EBNA-LP mRNAs containing the HF exon would not only destabilize these EBNA-LP mRNAs but also preclude the derivation of oligo-dT–primed cDNA reads containing both the HF exon and the EBNA-LP ORF. Therefore, our sequencing data likely greatly underestimate the percentage of EBNA-LP transcripts that contain the HF exon.

Supporting the role for pri-miR-BHRF1-2 and 1–3, but not pri-miR-BHRF1-1 in the *cis* regulation of EBNA-LP expression, previously published data demonstrate an upregulation of EBNA-LP expression in EBV with mutants individually lacking the pri-miR-BHRF1-2 or 1–3 stem-loop, but not with a mutant lacking only miR-BHRF1-1 (Feederle et al., 2011a). Additionally, the upregulation of the EBNA-LP-activated promoter Cp was only observed with pri-miR-BHRF1-2 and 1–3 single mutants, indicating that it is the upregulation of EBNA-LP expression that induces the observed increase in EBNA-LP-regulated transcription (Feederle et al., 2011a). We have used reporter assays to demonstrate that the pri-miR-BHRF1 and 1–3 stem-loops are indeed able to downregulate gene expression in *cis*, and we also demonstrate that this downregulation is, as expected, Drosha dependent (Fig. 5A). Moreover, we also observed a reduction in EBNA-LP protein expression when the HF exon was targeted using RNAi (Fig. 6). This demonstrates that destabilization of the HF exon, by cleavage by either Drosha or the RNA-induced silencing complex (RISC), both result in the inhibition of EBNA-LP expression.

While our data argue that the pri-miR-BHRF1-2 and 1–3 stem-loops are acting in *cis* to ensure that the optimal level of EBNA-LP protein in expressed, this does not, of course, indicate that the miR-BHRF1 miRNAs do not also regulate the expression of important cellular mRNAs in *trans*. However, these data do argue that the remarkable reduction in transformation potential and LCL growth rate seen with the ∆123 EBV mutant is primarily due to the overexpression of EBNALP, and/or its downstream viral gene targets, that is observed in the absence of these two viral pri-miRNAs. Furthermore, we identified two novel splice junctions (C2’ and W2-W1’ splicing), both of which would also reduce the expression of EBNA-LP. It therefore seems that EBV has evolved multiple mechanisms to post-transcriptionally downregulate EBNA-LP expression, further indicating that an optimal level of EBNA-LP expression is important for EBV pathogenesis.

We note that we do not address whether downregulation of EBNA-LP expression by RNAi, as shown in Fig. 6, increases the growth rate of the ∆123 LCLs, as predicted by our model (Fig. 8). This is because, even though we were able to stably express amiRNAs targeting the HF exon of EBNA-LP transcripts in∆123 LCLs, and this did reduce EBNA-LP expression by 2–5 fold (Fig. 6), we did not see a consistent increase in the growth rate of ∆123 LCLs that was statistically significant. This may in part reflect the ability of EBNA-LP to transcriptionally activate its own expression.

We note that the location of the miR-BHRF1-2 and 1–3 pri-miRNAs 3’ to both the BHRF1 and EBNA-LP ORF is evolutionarily conserved amongst primate lymphocryptoviruses (LCV) (Cai et al., 2006; Riley et al., 2010; Skalsky et al., 2014; Walz et al., 2010). Additionally, the EBNA-LP homologs in primate LCVs demonstrate identical functionality to EBV-encoded EBNA-LP (R Peng et al., 2000; Rongsheng Peng et al., 2000). Although the BHRF1-2 and 1–2* seed regions are conserved with other primate LCVs, mutating their seed sequence does not affect B cell transformation (Haar et al., 2015; Riley et al., 2010; Skalsky et al., 2014). Additionally, the seed sequence for miR-BHRF1-3 is not evolutionarily conserved amongst primate LCVs (Cai et al., 2006; Riley et al., 2010; Skalsky et al., 2014; Walz et al., 2010). Therefore, we hypothesize that the regulation of the expression of EBNA-LP-like proteins by viral pri-miRNA stem-loops acting in *cis* has been conserved across primate LCV evolution.

This research adds to a short list of mRNAs whose expression has been proposed to be downregulated by Drosha processing of a pri-miRNA-like stem-loop located in *cis*. In humans, transcripts encoding DGCR8, follistatin-like 1 (FSTL1), and the KSHV gene product Kaposin B (KapB) mRNAs are currently the only identified mRNAs that are regulated in this manner (Han et al., 2009; Lin and Sullivan, 2011; Sundaram et al., 2013). Indeed, a genome wide study of the human genome only found DGCR8 mRNAs to be downregulated by Drosha cleavage (Shenoy and Blelloch, 2009). However, there are at least two additional examples of mRNAs regulated by pri-miRNA stem-loops located in *cis* in mice: the transcription factors neurogenin 2 and T-box brain 1 (Chong et al., 2010; Knuckles et al., 2012; Marinaro et al., 2017). Interestingly, in most of these cases, the resulting miRNA is not thought to be expressed at a high enough level to be functional, unlike the KHSV and EBV miRNAs (Han et al., 2009; Knuckles et al., 2012; Marinaro et al., 2017; Sundaram et al., 2013). It therefore appears possible that KHSV and EBV have independently evolved taken advantage of both the *cis* and *trans* regulatory capabilities of miRNAs.

## Materials and Methods

### Cell culture

LCLs were generated from peripheral blood mononuclear cells (PBMCs) using both the WT and ∆123 EBV bacmids, as previously described (Feederle et al., 2011a, 2011b). LCLs and the EBV-negative Burkitt lymphoma cell line BJAB were grown in Roswell Park Memorial Institute (RPMI) 1640 medium supplemented with 10% fetal bovine serum (FBS), 50 μg/ml gentamicin (LifeTechnologies), and 1x Antibiotic-Antimycotic (Gibco, 15240062). 293T cells were grown in Dulbecco’s modified Eagle medium (DMEM) supplemented with 5% FBS, 50 μg/ml gentamicin, and 1x Antibiotic-Antimycotic. NoDice/∆PKR cells (Kennedy et al., 2015) and RNaseIII-/- 293T cells (Aguado et al., 2017) were grown in DMEM supplemented with 10% FBS, 50 μg/ml gentamicin, and 1x Antibiotic-Antimycotic. 293 EBV bacmid producer cells were grown in RPMI 1640 medium supplemented with 10% FBS, 50 μg/ml gentamicin, and 100 μg/ml hygromycin B (Corning). All cells were grown at 37°C with 5% CO_2_.

### Molecular clones

All miRNA and amiRNA expression vectors were generated by cloning into the pTREX vector using the XhoI and EcoRI sites present within the intron upstream of the Thy1.1 protein (primers 1–3 in Supplementary Table 1) (Poling et al., 2017). The pTREX BHRF1-123 miRNA expression vector was previously described (Poling et al., 2017). amiRNAs targeting the BHRF1 3’UTR, and a control FLuc amiRNA, were generated in the pri-miR-30 context, as previously described (Zeng et al., 2002), and then cloned into pTREX.

The BHRF1 3’UTR from both the WT and ∆123 EBV bacmids was PCR cloned into the 3’UTR of the pL-CMV-GL3 vector between the XhoI and NotI restriction sites (primers 4 and 5 in Supplementary Table 1) (Skalsky et al., 2012). The NanoLuc cDNA used as an internal control was cloned in place of FLuc, between NheI and XhoI restriction sites, to generate pL-CMV-NanoLuc (primers 6 and 7 in Supplementary Table 1) (Promega).

### Lentiviral transduction and selection

All lentiviral vectors were packaged in NoDice/∆PKR cells plated at 5 x 10^6^ cells per 15 cm dish and transfected the next day with 11 μg pCMV-∆R8.74, 4.4 μg pMD2.G, and 13.8 μg pTREX. Plasmids along with 73 μl polyethylenimine (PEI) were pre-mixed in 1 ml of OPTI-MEM (Gibco) then added to cells 15 min later. Cell culture media was changed the next day to RPMI supplemented with 10% FBS, 50 μg/ml gentamicin, and 1x Antibiotic-Antimycotic. Three days post-transfection, culture media were passed through a 0.45 µm filter and then used to transduce LCLs. Two days post-transduction, LCLs were treated with 0.2 μg/ml puromycin (puro). Puro levels were slowly increased over time until cells grew well at a final concentration of 0.8 μg/ml. LCLs were then induced with 1 μg/ml doxycycline (Dox) and subjected to flow cytometry two days later to test for Thy1.1 positivity using an APC-conjugated CD90.1/Thy1.1 antibody (BioLegend). Fully selected cells were used for cell based assays.

Lentiviral packaging plasmids pCMV-∆R8.74 and pMD2.G were gifts from Didier Trono, (Addgene plasmids #22036 & 12259). pCMV-∆R8.74 expresses all HIV-1 proteins except for *env*, *vpr*, *vif*, *vpu*, and *nef*. pMD2.G expresses the VSV-G envelope glycoprotein.

### Growth curve

Percent Thy1.1+ for pTREX vectors was determined using an APC-conjugated CD90.1/Thy1.1 antibody (BioLegend). LCLs were plated at 250,000 cells/ml in T25 flasks in 5 ml of media using 25% filtered spent WT LCL media and 75% fresh RPMI supplemented with FBS, 50 μg/ml gentamicin, 1x Antibiotic-Antimycotic and 1 μg/ml Dox, if transduced with pTREX. Cells were counted by flow cytometry on a FACSCanto II (BD) using AccuCount beads (Spherotech, Inc.) every 1–3 days and split back to 250,000 cells/ml. Cells transduced with pTREX were also tested for Thy1.1+. Cell counts were normalized to day 0 and then normalized to the WT LCL cell count. For pTREX transduced LCLs, only Thy1.1+ cells were considered for total cell counts.

### Stem-loop real-time PCR

miR-BHRF1 levels were determined by stem-loop real-time PCR as previously described (Chen et al., 2005; Feederle et al., 2011b; Poling et al., 2017). Briefly, total RNA was harvested from LCLs using TRIzol (LifeTechnologies) extraction following the manufacturer’s protocol except that the last 70% ethanol wash was replaced with a re-precipitation. Reverse transcription (RT) was performed using the TaqMan miRNA reverse transcription kit (Applied Biosystems), 10 ng total RNA, and stem-loop miRNA specific primers (ThermoFisher Scientific, Inc.). qPCR was then performed using 10 μl TaqMan universal PCR Master Mix No AmpErase UNG (Applied Biosystems), 4 μl of a 1:3 dilution of the 15 μl RT, 0.8 μl TaqMan miRNA specific probe, and 5.2 μl water on a StepOnePlus real-time PCR System (ThermoFisher Scientific, Inc.). U6 snRNA expression was analyzed alongside the miRNAs as an internal control. miRNA expression levels were normalized to WT expression, which was set to 1.

### Western blot

Cells were lysed in NP40 lysis buffer and sonicated. Protein amount was normalized using a Pierce BCA protein assay kit according to the manufacturer’s protocol (ThermoFisher Scientific). Expression of EBNA-LP, EBNA2, and LMP1 were probed with antibodies JF186, PE2, and S12 respectively. β-Actin (Santa Cruz sc-47778) was probed as a loading control. Band intensity was determined using GeneTools image analysis software (Syngene).

To identify knockdown of EBNA-LP protein by amiRNAs directed against the HF exon, ∆123 LCLs were transduced with pTREX-based vectors expressing the Luc amiRNA and HF exon targeting amiRNAs B4 and B11. Once fully selected with puro, as determined by Thy1.1 expression using an APC-conjugated antibody (BioLegend) and flow cytometry, cells were removed from puro media and induced with Dox. Four days post-induction, cells were harvested for Western blot.

### 4SU labeling

4-thiouridine (4SU) labeling was performed as previously described (Duffy et al., 2015). Briefly, 25 ml of LCLs were plated at 350,000 cells/ml. The next day, cells were treated with 200 uM 4SU for 1 h. Cells were then collected and total RNA was harvested with TRIzol following the manufacturer’s protocol except that the last 70% ethanol wash was replaced with an additional precipitation step. 50 μg total RNA at 1 μg/μl was combined with 5 μg MTS-Biotin-XX (Biotium) in 100 μl DMSO, 5 μl 1M Tris (pH 7.4), 1 μl 0.5 M EDTA, and 344 μl water. Tubes were covered with aluminum foil and allowed to rotate at room temperature for 30 min. Excess biotin was then removed and the RNA re-precipitated and brought up in 50 μl water. 10 μl were removed to be used as total input RNA. 40 μl MyOne Streptavidin C1 Dynabeads (ThermoFisher Scientific) were washed on a magnet three times with 0.1 M NaCl. Beads were resuspended in 40 μl 2x streptavidin binding buffer (10 mM Tris pH 7.4, 1 mM EDTA, 2 N NaCl). Beads were added to the remaining 40 μl RNA and allowed to rotate at room temperature for 30 min. Supernatant was then removed using a magnet and beads washed six times with wash buffer (100 mM Tris pH 7.4, 10 mM EDTA, 1 M NaCl). Nascent transcripts were then removed from the beads by 3x treatment with 100 μl 100 mM dithiothreitol (DTT). Nascent RNA was then reprecipitated and resuspended in 20 μl water. Nascent and total RNA were reverse transcribed using the High-Capacity cDNA Reverse transcription kit according to the manufacturer’s protocol using 2 μg total RNA and 2 out of the 20 μl of nascent RNA (ThermoFisher Scientific). cDNA was diluted with 100 μl to make 120 μl total and 3 μl was used per qPCR reaction. qPCR was performed with Power SYBR Green Master Mix (ThermoFisher Scientific) for WT and ∆123 total and nascent transcripts using the primers 11–20 in Supplementary Table 1. Half-life was calculated using the formula described by Dolken et al. (Dölken et al., 2008). Nascent transcription was determined by the ∆∆Ct method normalizing all nascent transcript expression first to GAPDH and then normalizing ∆123 nascent expression to WT, which was set to 1. Total transcription levels were determined the same way except using total transcript expression levels.

### RNA-seq

Quality-controlled paired-end fastq files were pulled from ten random donors from the 1000 genomes project (Lappalainen et al., 2013). Reads were first aligned to the human genome (hg19) using HISAT2 (Kim et al., 2016). Unaligned reads were aligned to the WT EBV genome (Accession: NC_007605). Data were visualized in IGV and splice junctions read depth was quantified (Robinson et al., 2011).

### PacBio sequencing

PacBio sequencing was performed by isolating Cp/Wp transcripts from total B95-8 LCL RNA using a W2 3’-biotinylated oligonucleotide (primer 21 in Supplementary Table 1). Briefly, 50 μg total RNA was mixed with 250 pmol of the W2 probe. RNA and probe were heated to 95°C for 5 min and then allowed to slowly cool to room temperature. 20 μl MyOne Streptavidin C1 Dynabeads (ThermoFisher Scientific) beads washed and resuspended in 2x streptavidin binding buffer as described above. RNA annealed to probe and beads were mixed and rotated at room temperature for 30 min. Beads were washed with wash buffer as described in section 3.5.8. Isolated transcripts were released from the beads by resuspending the beads in 100 μl wash buffer, heating to 95°C for 5 min and immediately adding to a magnet and collecting the supernatant. Isolated transcripts were then reprecipitated and resuspended in 20 μl water. Transcripts were then subject to an Iso-seq cDNA library preparation using the SMARTer PCR cDNA synthesis kit (Clontech) and size selected for 800 - 2 kb and 2 kb+ sized transcripts. SMRTbell adapters were then added to the isolated libraries and samples were sequenced on a PacBio RS II. Library preparation and sequencing was performed by the Duke sequencing and genomic technologies shared resource core. Sequence data were then aligned to the human genome (hg19) using GMAP (Wu and Watanabe, 2005). Unaligned sequences were then aligned to the B95-8 EBV genome (Accession: V01555).

### Luciferase assay and Northern blot

293T cells and RNaseIII-/- cells were plated at 1x10^5^ cells per well in a 24-well plate for both luciferase assay and Northern blot. The next day cells were transfected with 1 ng pL-CMV-NanoLuc, 10 ng pL-CMV-GL3, pL-CMV-GL3 BHRF1-3’UTR, or pL-CMV-GL3 BHRF1-3’UTR ∆23, and 500 ng pcDNA3 as a filler along with 1.27 μl polyethylenimine (PEI). Media was changed the next day and two days post-transfection cells were harvested with 1x passive lysis buffer (Promega). Firefly and NanoLuc expression was determined using the Nano-Glo dual-luciferase reporter assay system (Promega). Firefly expression was normalized to NanoLuc expression. pL-CMV-GL3 BHRF1-3’UTR and pL-CMV-GL3 BHRF1-3’UTR ∆23 relative expression levels were normalized to the pL-CMV-GL3 relative expression level. Last, the 293T transfected cells expression levels were normalized to the RNaseIII-/- expression levels.

Two days post-transduction total RNA was isolated by TRIzol extraction as described above. 10 μg total RNA was run on a 15% TBE-Urea gel (BioRad). RNA was transferred onto nylon membrane (Perkin Elmer) which was dried out and crosslinked (Stratalinker, Stratagene). The membrane was then treated with express-hyb hybridization solution (Clontech) for 1 h at 37°C. U6 and BHRF1-2, 1–2*, and 1–3 complementary oligos were labeled with gamma-p32 ATP using T4 PNK (NEB) (primers 22–25 in Supplementary Table 1). Labeled oligos were allowed to incubate with the nylon membrane for 1 h at 37°C and then washed. Membranes were exposed to Biomax MS film.

### Shallow sequencing

EBNA-LP shallow sequencing was performed by first generating cDNA from total B95-8 LCL RNA using SuperScript IV according to the manufacturer’s protocol (ThermoFisher Scientific). EBNA-LP transcripts were then PCR amplified and cloned into the SpeI and PstI digestion sites in pGEM-5ZF+ (primers 26 and 27 in Supplementary Table 1) (Promega). Individual clones were isolated and sequenced using the T7 and SP6 sequencing primers.

## Acknowledgments

The research reported in this manuscript was supported by National Institute of Health grant R01-AI067968. B.C.P. and A.M.P. were supported by T32-CA009111 and A.M.P. was also supported by F31-CA180451.

The authors thank Benjamin tenOever and Didier Trono for the gift of reagents used in this research. The authors would also like to thank the Duke Cancer Institute Flow Cytometry Shared Resource Core for their help with collecting flow cytometry data. Lastly, the authors would like to thank the Duke sequencing and genomic technologies shared resource center, especially Olivier Fedrigo, for help with the PacBio sequencing.

---

**Supplementary Table 1.**
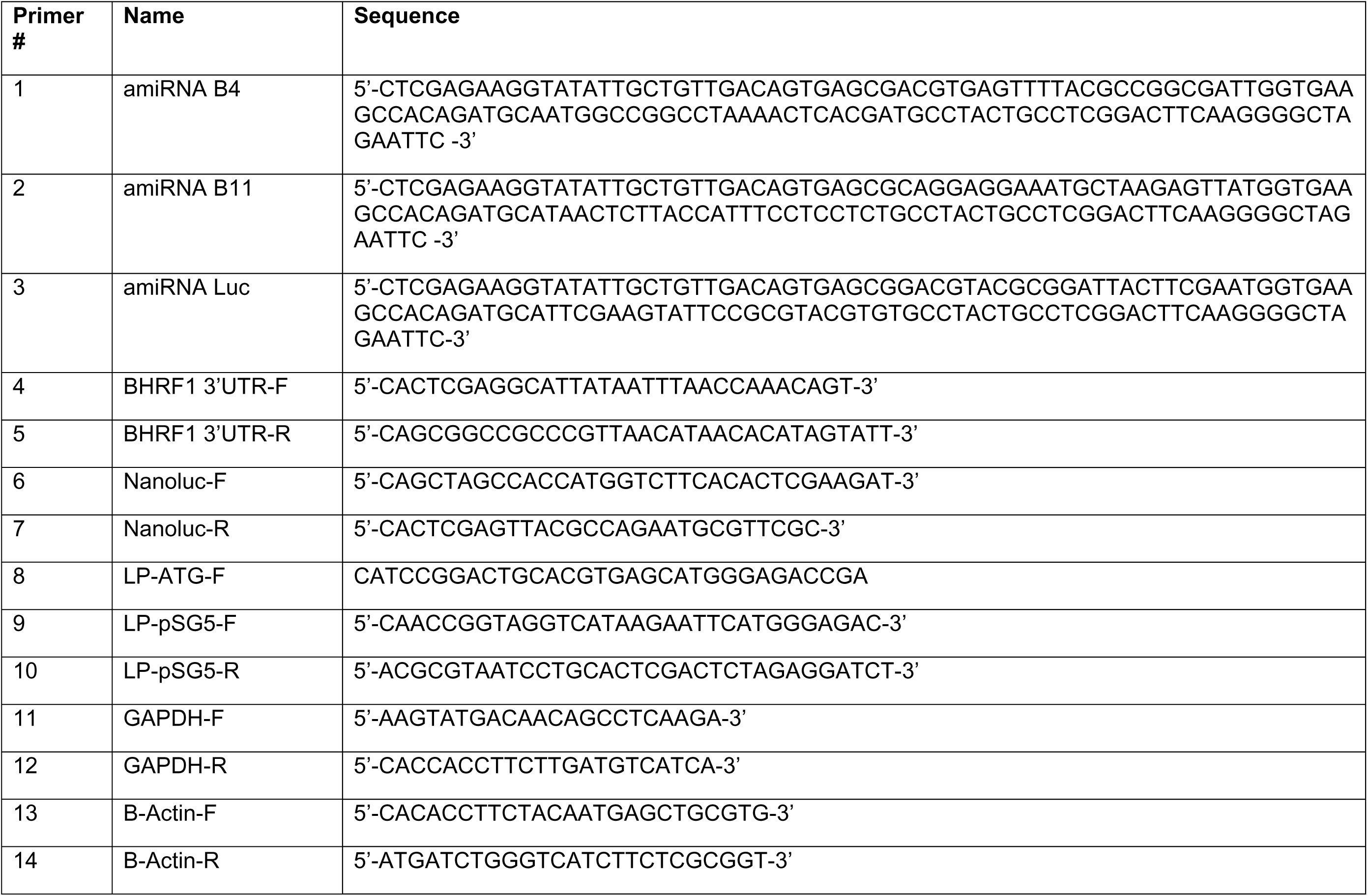
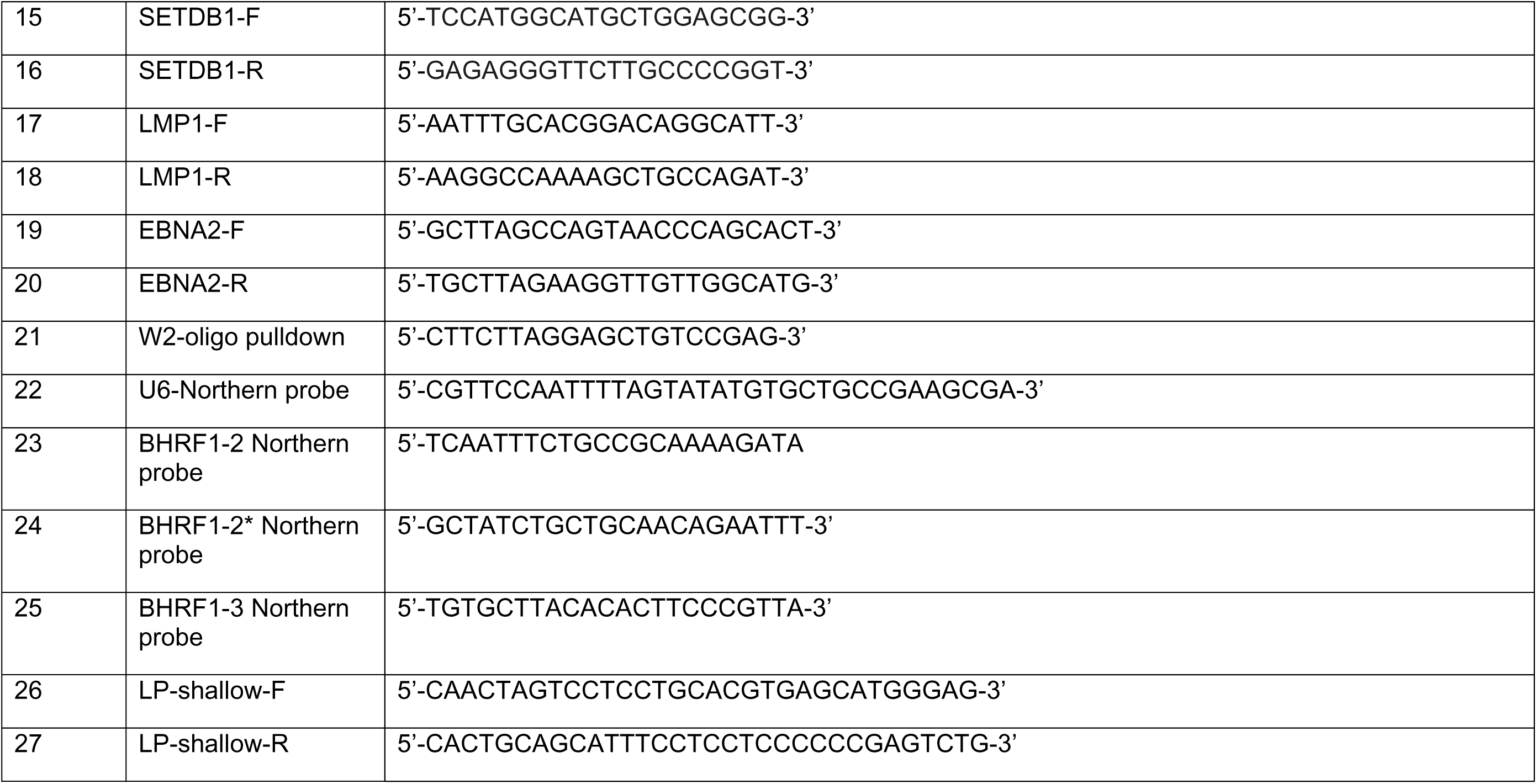
List of relevant sequences and primers.

